# Predicting clinical phage therapy outcomes *in vitro*: results using mixed versus single isolates from an MRSA case study

**DOI:** 10.1101/2025.04.28.650979

**Authors:** Meaghan Castledine, Chloe Esom, Brieuc Van Nieuwenhuyse, Sarah Djebara, Maya Merabishvili, Jean-Paul Pirnay, Angus Buckling

**Affiliations:** Centre for Ecology & Conservation, University of Exeter, Penryn Campus, Penryn, TR10 9FE; Institute of Experimental and Clinical Research, Université catholique de Louvain - UCLouvain, Brussels, Belgium; Center for Infectious Diseases, Queen Astrid Military Hospital, Brussels, Belgium; Laboratory for Molecular and Cellular Technology, Queen Astrid Military Hospital, Brussels, Belgium, 1120

**Keywords:** bacteriophages, applied microbiology, clinical microbiology, staphylococcus, microbial ecology

## Abstract

**Aims:** In phage therapy case studies, 1-3 bacteria isolates are typically tested against phages (phagogram). However, as bacteria populations differ in their susceptibility to phages and antibiotics, the strains selected may not represent how the infecting population will respond to treatment. Our aim was to assess whether the effects of phage on single or a mix of isolates *in vitro* show more comparable results to that observed during a clinical case study.

**Methods and Results:** The patient presented with a methicillin resistant *Staphylococcus aureus* infection (MRSA). In this previously published case study, phage therapy alongside antibiotics rapidly cleared blood cultures of bacteria while localised regions, including the lungs, took longer to clear of bacteria. In this follow-up study, mixed isolates were more likely to persist than single isolates *in vitro*, more closely representing the lung, but not blood, infections. These results may reflect the different degrees of genetic diversity of the infecting bacteria in these sites.

**Conclusions:** For this patient, phage therapy predictions were significantly affected by whether we used mixed versus single isolates, but the predictive precision depended on the site of *in vivo* infection.

**Impact statement:** This work is important for informing whether phage therapy design protocols give accurate insights into clinical outcomes.

## Introduction

4.95 million deaths were associated with antimicrobial resistant (AMR) infections in 2019, with this figure expected to rise to 8.22 million by 2050 (WHO 2019; Naghavi et al. 2024). AMR is driving a need for complementary therapeutics such as phage therapy (Abedon 2019; Pirnay et al. 2024) which uses the viruses (bacteriophages; phages) that infect and kill bacteria to treat bacterial infections. Phages are advantageous as phage resistance can be physiologically costly to growth, often resulting in resistant populations becoming more susceptible to the immune system or antibiotics (Oechslin et al. 2016; Roach et al. 2017; Oechslin 2018). Phages are also ubiquitous and diverse, facilitating ease of discovery and relatively cost-effective to manufacture (Abedon 2019).

Since bacteriophages are host-specific, phages are commonly selected for treatment on a case-by-case basis to ensure high efficacy against the infecting bacterium (Schooley et al. 2017; Abedon et al. 2021; Köhler et al. 2023; Pirnay et al. 2024). When designing patient-specific therapeutics, typically 1-3 bacteria isolates (Sarker et al. 2016; Schooley et al. 2017; Chan et al. 2018; Dedrick et al. 2019; Nir-Paz et al. 2019; Lebeaux et al. 2021; Eskenazi et al. 2022; Simner et al. 2022; Blasco et al. 2023; Otava et al. 2024) are taken from the patient and screened against a panel of phages for infectivity (‘phagograms’) (Kolenda et al. 2024). The phages which are the most effective at preventing bacterial growth are selected for treatment. *In vitro* experiments provide a valuable tool for predicting treatment outcomes (Castledine et al. 2022), and designing treatments to optimise success for individual patients (Schooley et al. 2017; Pirnay et al. 2024).

However, an issue with using a small number of bacteria isolates for phagograms is population heterogeneity. Bacteria populations are genetically diverse and multiple lineages can coexist within one infecting population (Klemm et al. 2016; Didelot et al. 2016; Launay et al. 2021; Hoang et al. 2024). Long-term studies of cystic fibrosis patients have shown population diversification within chronic infections (Mowat et al. 2011; Lieberman et al. 2014). These strains can vary in their virulence, antibiotic and phage sensitivity (Mowat et al. 2011; Lieberman et al. 2014; Didelot et al. 2016; Castledine et al. 2022). For instance, in a recent case study, 1/24 *Pseudomonas aeruginosa* isolates was resistant to one phage (*Pseudomonas aeruginosa* bacteriophage: 14-1) in the cocktail while susceptible to the other (*Pseudomonas aeruginosa* bacteriophage: PNM), with the remaining 23/24 isolates being completely susceptible to both phage (Castledine et al. 2022). The rapid diversification of bacteria, owing to their high mutation rates (Jayaraman 2011) also means populations may change between sending an isolate for a phagogram and starting phage therapy. Consequently, by chance a bacterium isolate may be selected for phagograms that does not accurately capture how the whole population will respond to phage therapy.

This project builds from a recent phage therapy case study in which a 12-year old boy was treated with bacteriophage ISP for an *Staphylococcus aureus* infection (Van Nieuwenhuyse et al. 2024). The *S. aureus* was methicillin resistant (MRSA) and panton-Valentine leukocidin-producing which is a toxin specifically adapted to killing myeloid (immune) cells. The infection was treated successfully with a combination of phage therapy and antibiotics (Van Nieuwenhuyse et al. 2024). From a sample taken on the same day as phage therapy started, we obtained 24 *S. aureus* isolates. We compared the efficacy of ISP, and resistance evolution, against 4 individual isolates and a mixture of all 24 isolates *in vitro*. 24 isolates were selected as a feasible number for phage therapy centres to mix for phage therapy predictions, therefore capturing as much diversity as feasibly possible. This may include multiple isolates of the same strain, however this will be more representative of the total bacterial population that when using single isolates. We further compared the *in vitro* results to those found *in vivo* during phage therapy.

## Methods

### Ethics statement

The study was performed in accordance with the ethical standards as laid down in the Declaration of Helsinki and as revised in 2013. The patients gave informed consent and their anonymity was preserved. Details of ethical approval for phage therapy can be found in the corresponding case study (Van Nieuwenhuyse et al. 2024).

### Patient information and sample extraction

The 12-year-old male patient presented with Panton-Valentine Leukocidin-producing methicillin resistant *Staphylococcus aureus* (MRSA). Full details of his treatment are published (Van Nieuwenhuyse et al. 2024), and key details are summarised here. Infection started from a scratch coinciding with minor ankle trauma and progressed to generalized necrotizing fasciitis, and vascularly to his lungs and heart. Phage therapy using a single bacteriophage, ISP, was started nine days following hospital admission and continued daily via multiple routes for a total of 22 days. ISP infects via glycosylated wall teichoic acids (WTAs) and the WTA backbone on the cell surface. ISP has been used in numerous phage therapy cases as a broad host-range *Staphylococcus aureus* phage (Merabishvili et al. 2009; Xia et al. 2011; Takeuchi et al. 2016; Pirnay et al. 2024).

For this study, two samples were obtained at the start of treatment, one of pus and one of pleural fluid. By comparing bacteria from the start and end of treatment, we can determine if there are any changes in phage resistance. Infection and treatment progress in the patient was not determined from these samples with respect to informing treatment practices – these samples are for retrospective analyses on resistance evolution. Three samples were taken after six days of treatment including two pleural fluids (one taken pre-phage dose and one after phage dose) and one bronchoalveolar lavage (BAL). Samples were plated directly onto lysogeny broth (LB) agar and 24 bacteria colonies were isolated per sample (Table S1). Individual isolates were grown in 150 μL LB media overnight at 37°C, then 150 μL 50% glycerol was added and samples frozen at -70°C. No bacteria were detected in pleural fluid samples from day six. Bacteriophages were isolated via chloroform extraction: 900 μL of sample was vortexted with 100 μL chloroform then centrifuged at 14000 rpm (21100 *g*) for five minutes and the supernatant isolated. Supernatant (10 μL) was then added to 6 mL LB media with 60 μL *S. aureus* PS overnight culture (patient strain originally isolated for phage testing) and grown overnight from which phages were extracted (via chloroform extraction). Amplification of extractions was conducted to purify phages from contaminants including antibiotics. Pleural fluid from day six were the only samples which contained phage (Table S1). Amplified phage stocks were diluted 100-fold and inoculated into LB soft agar with 30 μL *S. aureus* PS (from overnight culture) which was poured onto an LB agar plate. Plates were grown overnight at 37°C. Phage plaques were picked (24 per phage population) and grown individually with 60 μL *S. aureus* PS. Phages were extracted as previous and pooled into their respective populations (i.e. 24 isolates within each of the two populations). In summary, bacteria were isolated from two samples at the start of treatment and one sample at day six of treatment. Phages were isolated from two samples at day six of treatment.

### *In vivo* bacteriophage resistance

Bacteria from the start (n = 48 isolates, 24 / sample) and day six of treatment (isolated from lungs; n = 24) were tested for resistance to ISP and phage isolated from day six (two samples). Isolates were grown overnight in 180 μL LB media. Cultures were diluted to 0.2 OD_600_ with 200 μL added to 12 mL LB soft agar and poured onto square LB agar plates. 5 μL dilutions of phage stocks from undiluted to 10^-8^ dilution were spotted onto lawns of each host. Phage resistance was estimated by the relative ability of each phage to plaque onto each host strain (lower phage densities estimated on more resistant hosts).

### *In vitro* experimental evolution

We next considered how phage therapy predictions made using laboratory experiments differed depending on whether single or mixed-isolate cultures were used. Treatments included four individual isolates, and a master-mix of 24 isolates (how the master-mix is generated is detailed below) cultured in the presence and absence of phage for six days (Figure 1). Each treatment (phage present / absent) for each individual isolate was replicated 6 times (e.g. 6 cultures of isolate 1 with phage and 6 cultures of isolate 1 without phage). The mixed-isolate treatments were replicated 12 times.

**Figure 1.**
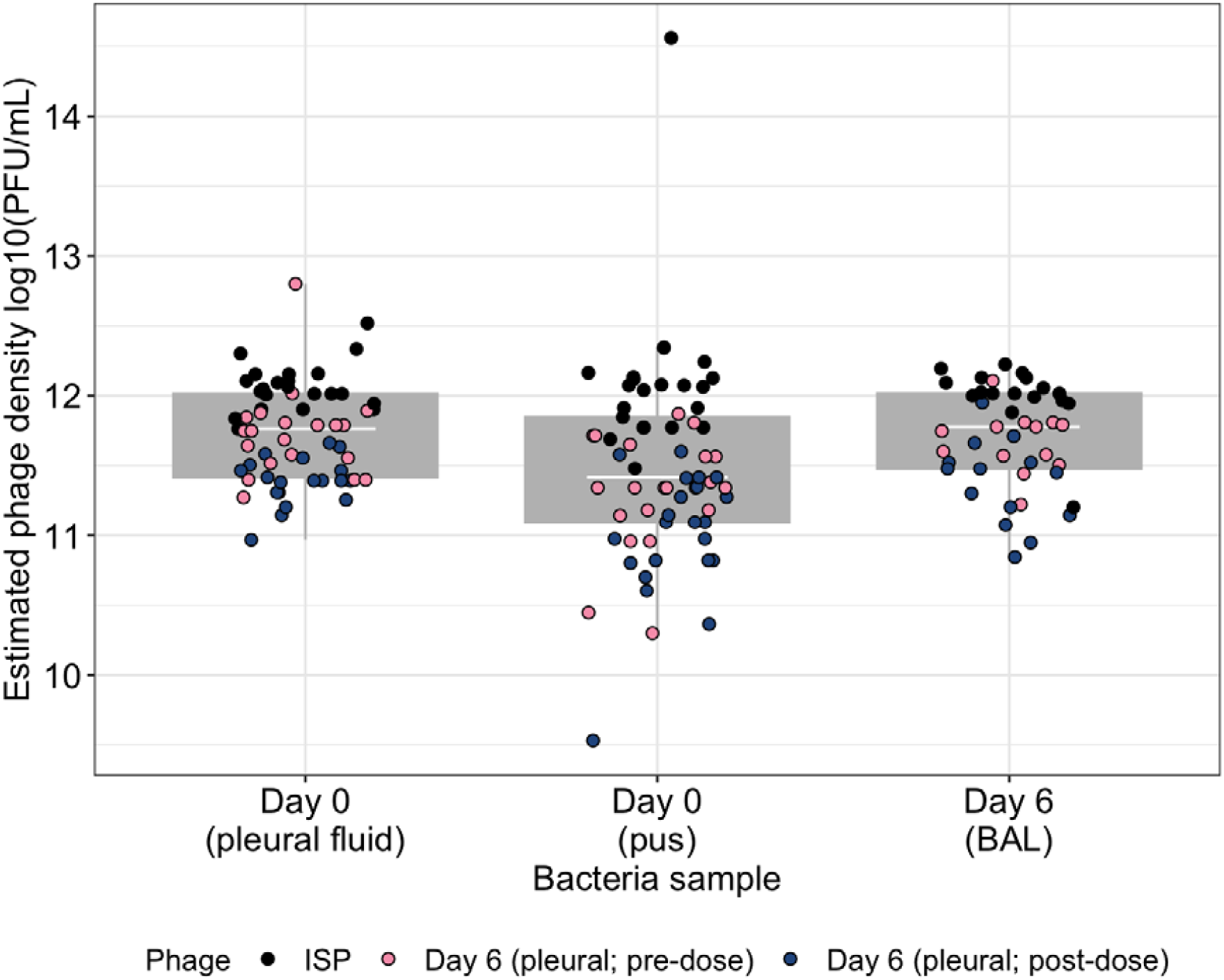
Phage densities of ISP and two day 6 phage stocks when plated onto lawns of *S. aureus* isolates isolated from day 0 (start of treatment) and day 6 of treatment (two samples, one before and one after treatment). High plating densities of all phages indicate a lack of resistance evolution during phage therapy treatment. Individual points represent individual isolates phages were plated on isolated from the indicated time. Tops and bottoms of the bars represent the 75th and 25th percentiles of the data, the middle lines are the medians, and the whiskers extend from their respective hinge to the smallest or largest value no further than 1.5* interquartile range.

The master-mix of day 0 isolates (from pleural fluid) was generated by growing the 24 isolates overnight, then normalising densities to 0.1 OD_600_ and mixed at equal ratio. 900 μL master-mix was frozen with 900 μL 50% glycerol and stored at -70°C. Four isolates from day 0 (pleural fluid) were picked at random. Individual isolates and the master-mix (60 μL) was grown overnight in 6 mL LB media. Cultures were diluted to 0.1 OD_600_ (2.4 x 10^6^ CFUs / μL) and 60 μL (144 x 10^6^ CFUs) were inoculated into relevant vials containing 6 mL LB media. To ‘phage present’ cultures 21.8 μL of phage stock (6.6 x 10^5^ PFU/μL) was added for an MOI of 0.1. Cultures were incubated at 37°C and shaken at 180 rpm. Every 24 hrs, cultures were vortexed and 60 μL transferred to fresh LB media. To ‘phage present’ cultures, 21.8 μL of phage stock was inoculated at each transfer. Each transfer, samples were frozen (as above, master-mix) and phages extracted via chloroform extraction as previous. Bacterial densities were estimated by plating cultures, from frozen, onto LB agar. Phage densities were estimated by plating phages onto lawns of *S. aureus* strain 13 S44 S9.

### *In vitro* bacteriophage resistance

Bacteriophage resistance was estimated from day 6 *in vitro* cultures. 12 isolates were picked from each treatment replicate and grown overnight in 200 μL LB media. Phage resistance was estimated via spot tests of ISP onto lawns of each isolate as previous. Isolates were scores as 1, susceptible (clear zone of lysis) or 0, resistant (no growth inhibition).

### Statistical analyses

All data were analysed using R (v.4.2.1) in RStudio (Team 2013) and all plots were made using the package ‘*ggplot2*’ (Wickham 2016). Model simplification was conducted using likelihood ratio tests and Tukey’s post hoc multiple comparison tests were done using the R package ‘*emmeans*’ (Lenth 2018).

Estimates of *in vivo* susceptibility to bacteriophage ISP and *in vivo* phages was confounded by a block effect of ‘day’ samples were tested with samples in the 2^nd^ block having 29.8-41.7% reduction in phage titres across all bacteria-phage combinations. A correction was applied to estimates in the second block accounting for the average difference in each bacteria-phage combination (e.g. day 0 pus bacteria vs ISP, day 0 pus bacteria vs day six pre-dose phage).

Changes in bacteria density in vitro were estimated using a linear mixed effects model. Here, bacteria densities (log10 CFU/mL) were analysed against interacting fixed effects of treatment (isolate 1-4 or mixed isolate), phage presence and time (day of transfer) with a random effect of treatment replicate. Phage densities (log 10 PFU/mL) were analysed in a linear mixed effects model with phage density analysed against interacting fixed effects of treatment and time with a random effect of treatment replicate. To assess whether phages were replicating above baseline inoculum (the volume of phage inoculated at each transfer), phage densities were analysed in multiple one-sample t-tests. If phages were not significantly replicating, densities would be non-significantly different to 6.38 log10 PFU/mL. P-values were corrected for multiple testing using the ‘false-discovery-rate’ method.

## Results

### No evidence of phage resistance *in vivo*

The patient’s treatment followed initially favourable responses to phage therapy, with clearance of blood cultures (bacterial extinction) within 30 hours while bacteria persisted in the lungs for at least six days; these bacteria were successfully treated with a combination of phage and antibiotics (Van Nieuwenhuyse et al. 2024). The persistence of bacteria in the lungs six days into phage therapy (last sample taken) presents the possibility of phage resistance emergence which may have delayed bacteria clearance. Consequently, we tested whether any bacteria isolated from the start or day six of treatment (isolated from lungs) were phage resistant.

Bacteriophage stocks of ISP and two phage populations isolated (and amplified) from day six of treatment (one before and one following phage dose) were plated on different bacteria isolates isolated at the start (day 0) and end (day six) of treatment. Evolution of phage resistance would be evident by a phage plating densities being lower on day six bacteria compared to day 0 bacteria. However, all phages plaqued onto bacteria hosts with very high efficiency (ISP: 11.2 – 12.5 (one outlier at 14.6) log10(PFU/mL); day six pre-dose: 10.3 – 12.8 log10(PFU/mL); day six post-dose: 9.5 – 11.9) suggesting a lack of resistance to any phage isolates (Figure 1). Differences in plating densities between phages is reflective of their stock densities rather than changed in infectivity. As there is no evidence of phage resistance evolution, we would not expect significant changes in phage infectivity.

### *In vitro* population dynamics

We next considered if cultures composed of a mixed population (24 isolates) or single isolates differed in their likelihood of persistence when exposed to ISP. Extinction rates were markedly higher in single isolate populations (58.3% extinction rate across single isolate replicate populations; Figure 2). All replicates of isolate 4 were extinct by day two, while 3/6 replicates of isolates 1 and 3 went extinct, and 2/6 of isolate 2 went extinct by day 6. Comparatively, only 1/12 (8.3%) replicates of the mixed-isolate population went extinct. Differences in extinction rates led to mixed-isolate populations having an significantly higher densities (X = 4.56, 95%CI = 3.78 – 3.55) when infected with phage compared to 3 out of 4 single isolate populations (isolate 1: X = 1.62, 95%CI = 0.53 - 2.71; isolate 3: X = 1.61, 95%CI = 0.51 - 2.70; isolate 4: X = 0.15, 95%CI = -0.94 - 1.24; ANOVA comparing models with and without treatment x phage interaction: X^2^ = 92.75, p < 0.001; Table S2). Only one isolate (isolate 2) was non-significantly different to the mixed-population (X = 2.66, 95%CI = 1.56 - 3.76; Tukey HSD: estimate = -1.89, t-ratio = -2.840, p = 0.051; Figure 2). Differences between mixed and single-isolate populations were not explained by differences in ability to reach higher densities in general, with no significant differences between no-phage controls over time (Figure 2; Table S2). These different patterns observed between different single isolates, and between single to mixed-isolate populations demonstrate phenotypic diversity in how each individually and collectively respond to phage infection. This phenotypic diversity is presumably reflective of genetic diversity which may be in genetic mutations, carriage of different mobile genetic elements, or phase variants (different genes turned ‘on’ or ‘off’).

**Figure 2.**
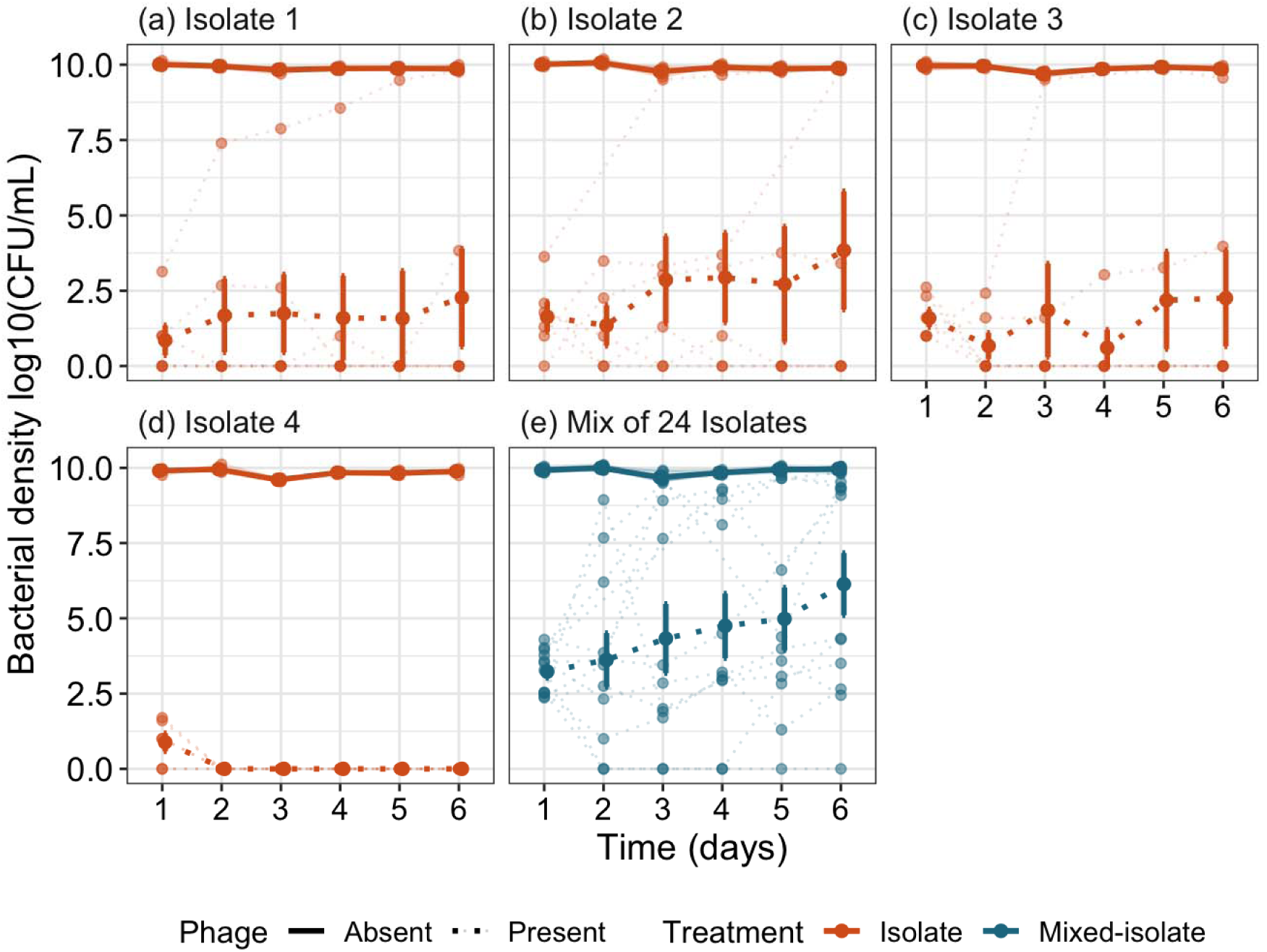
Bacteria densities when grown in the presence and absence of phage for 6 days. Bacteria populations started with either a single *S. aureus* isolate isolated from day 0 of phage therapy (a-d) or a mix of 24 *S. aureus* isolates isolated from day 0 (e). Results indicated higher extinction rates when experiments were started with single isolates, compared to mixed isolates which had higher rates of persistence at low density and recovery to densities dissimilar to controls. Solid points and bars represent the treatment mean and standard error (SE). Faint points show the treatment replicates with lines connecting the same replicates through time.

Of the replicates which ‘survived’, populations either persisted at low density (X = 3.56 log10(CFU/mL) ± 0.25 SE (standard error)) or recovered (X = 9.54, ± 0.085 SE) with densities comparable to no-phage controls (X = 9.9, ± 0.013 SE). Mixed populations had a higher rate of recovery with 6/12 (50%) replicates compared to 4/24 (16.7%) replicates of single isolate populations. Population recovery led to bacteria populations having significantly greater density at the end vs start of the experiment (ANOVA comparing models with and without phage x time: X^2^ = 12.64, p = 0.027; Table S3). Persistence was also higher in mixed isolate populations with 5/12 (41.7%) populations while 3/24 (12.5%) of single isolate populations persisted at low density. Consequently, when examining effects of phage between mixed and single isolate populations, the single isolates were more likely to predict complete extinction while mixed populations were more likely to predict population persistence and recovery with phage. Extinction predictions using single isolates reflected the initial blood results found *in vivo* with clearance of bacteria, while predictions using mixed isolates are more reflective of localised persistence of bacteria within the lungs.

Phages replicated on day 1 of the experiment above the base-line inoculum in each treatment (Figure 3; Table S4). From days 2-6, phage populations did not increase above inoculum levels in most treatments (Figure 4; Table S4). However, there was considerate variation, particularly in the mixed-isolate population, which will be discussed further in the context of phage resistance and bacteria population density.

**Figure 3.**
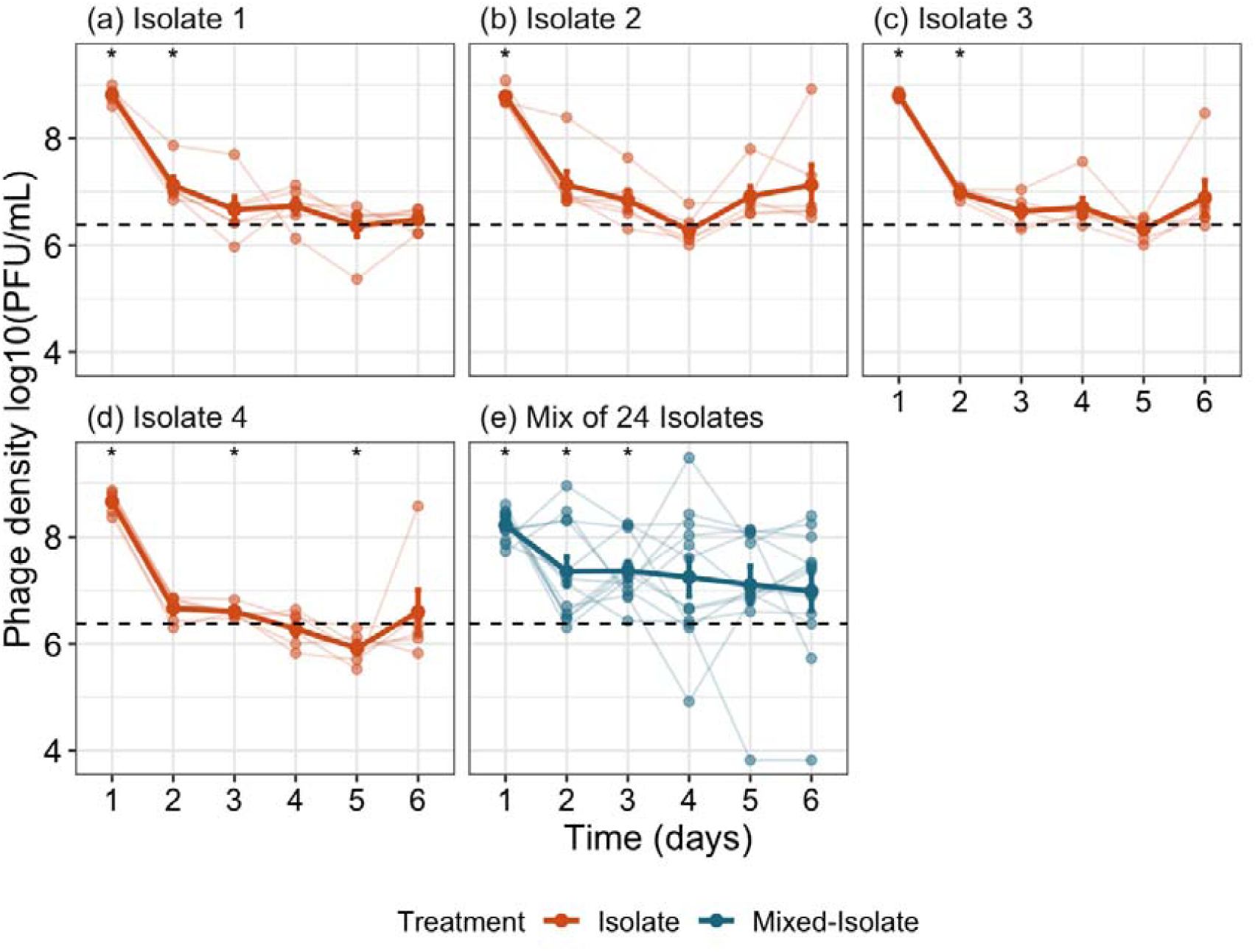
Phage densities through time between bacteria cultures initiated with individual isolates (a-d) and mixed-isolates (e). Phages had high levels of replication on day 1 with densities reducing to inoculum levels across the experiment, although this decline was slower where experiments were initiated with a mix of clones. The dashed line indicates the density of phage inoculated at each transfer while asterisks (*) indicate where phage densities during experiment are significantly different from that value. The solid points and bars represent the treatment mean and standard error (SE). Faint points show the treatment replicates with lines connecting the same replicates through time.

**Figure 4.**
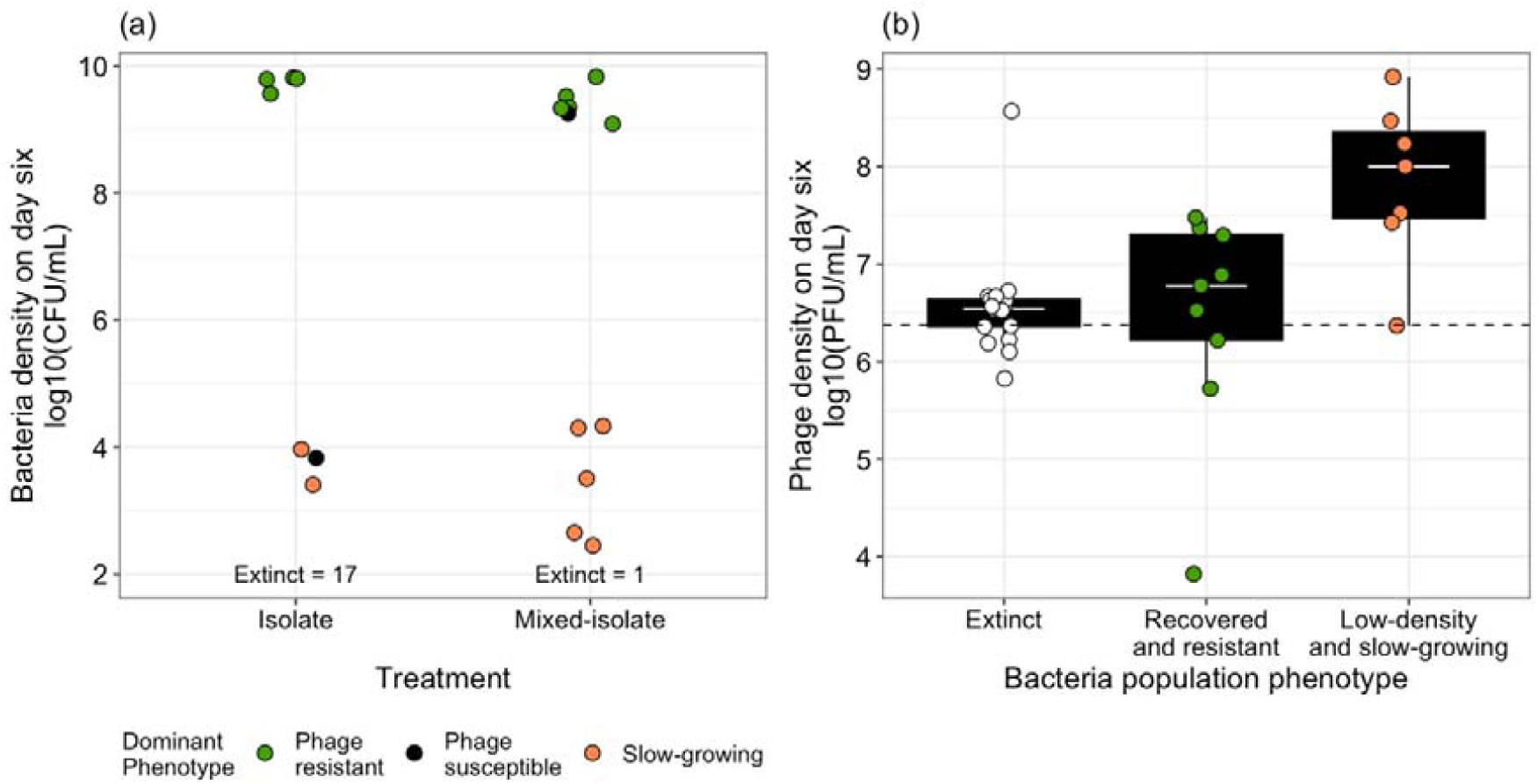
(a) Associations between bacteria resistance and density at day six, demonstrating that population recovery (high densities) are associated with phage resistance, while low-density populations have colonies which are too slow-growing for resistance testing. (b) In the low-density populations, phages are shown to be replicating above base-line levels implying a lack of complete phage resistance. In contrast, phages are not replicating in the recovered/phage resistant population, as comparable to where bacteria are extinct, demonstrating that phage resistance is inhibiting phage replication in these cultures. Points represent individual treatment replicate populations. Tops and bottoms of the bars represent the 75th and 25th percentiles of the data, the middle lines are the medians, and the whiskers extend from their respective hinge to the smallest or largest value no further than 1.5* interquartile range.

### Phage resistance *in vitro*

*In vivo*, we found no evidence of phage resistance and we next tested whether phage resistance emerged *in vitro* within persisting or recovered populations. Populations refer to separate experimental replicates of each single-isolate or mixed-isolate treatment. From these replicate populations, 12 colonies were tested for phage resistance – therefore each replicate population is composed of 12 colonies.

4/24 single isolate replicate populations and 6/12 mixed-isolate replicate populations recovered in density comparable to controls. In all 4 ‘recovered’ single isolate populations, all colonies (12/12) tested were phage resistant. In 4/6 recovered mixed-isolate replicate populations, all colonies (12/12) were phage resistant while 11/12 colonies in one replicate population were resistant (the remaining colony was slow-growing – see below on slow-growing colonies); and 1/6 replicate recovered mixed-isolate populations was completely phage susceptible (0/12 resistant). These results suggest that populations recovered via fixation of phage resistance (Figure 4a). This is further demonstrated in the day six phage-density data (from resistant cultures) which shows phage populations not replicating above inoculum levels (one sample t-test, t-value = 0.201, p_fdr_ = 0.846; Figure 4b). This is comparable to phage densities in extinct-bacteria populations which show no difference above base-line inoculum (one-sample t-test, t-value = 1.45, p_fdr_ = 0.248; Figure 4b).

Low-density populations were dominated by colonies which could not be cultured to densities high enough for phage-resistance testing, suggesting the low-density of the populations were due to very slow growth rates (Figure 4a). These could have been present *in vivo* and contributed to treatment challenges but remained undetected if sampling was biased to faster-growing populations. 3/24 single isolate replicate populations and 5/12 mixed-isolate replicate populations were found to have low-densities (X = 3.56 log10(CFU/mL) ± 0.25 SE (standard error)). In two low-density single isolate populations, 1 or 2 colonies were phage resistant while the remaining were slow-growing. In the 3^rd^ single isolate population, 8/12 colonies were phage susceptible while 4/12 were slow growing. All colonies from low-density mixed-isolate populations were slow-growing. A lack of complete phage resistance in these slow-growing populations is implied by phage densities being significantly higher than inoculum density (one sample t-test, t-value = 4.66, p_fdr_ = 0.01; Figure 4b), suggesting phages were replicating. Phage replication would further contribute to diminished bacteria densities in these low-density populations. The slow-growing colonies were also, observationally, smaller than resistant or susceptible colonies when grown on agar. This potentially suggests these colonies had slower growth rates which is why colonies did not grow enough in media for resistance testing. Slowed bacteria growth rates can reduce phage infectivity, allowing bacteria to persist. As we observed slow growth of colonies in the absence of phage, and had measurable phage replication during experimental evolution, low cell densities during the experiment were likely a product of slow growth and cell lysis.

## Discussion

We analysed a phage therapy case study to examine whether treatment outcomes were better predicted *in vitro* using mixed rather than single isolate populations. This case study was a complex MRSA infection in which phage therapy and antibiotics successfully cleared blood of *S. aureus* within 30 hrs while persisting for at least six days in the lungs (Van Nieuwenhuyse et al. 2024). Neither single or mixed-isolate populations completely captured *in vivo* dynamics. Single isolate populations showed high rates of extinction, comparable to results showing clear blood cultures within 30 hrs of treatment, while overestimating complete bacteria eradication. In contrast, mixed-isolate populations showed higher rates of bacteria persistence, reflecting *in vivo* persistence in lung tissue. However, mixed-isolate populations were more likely to increase in density with fixation of phage resistance than single isolate populations, while no resistance was observed *in vivo*. These results highlight limitations for predicting treatment outcomes using *in vitro* experiments, particularly for highly complex cases that are likely influenced by multiple variables, including host immunity, antibiotics and co-occurring microbes.

The differing abilities of single versus mixed *in vitro* populations to capture *in vivo* dynamics may be a consequence of different levels of diversity in the blood versus lung infections. The isolates were obtained from the lungs and it was this *in vivo* environment that was better captured by mixed versus single isolates. The lung environment is relatively more heterogenous than the bloodstream (Lieberman et al. 2014), and this heterogeneity can support a greater genetic diversity within the bacterial population (Brockhurst et al., 2004). In this patient’s case, he had an endotracheal tube which would have further increased lung heterogeneity and this was reflected by the empyema (collection of pus) within the lungs; this likely contributed to delayed treatment success with any biofilm and pus offering bacteria a ‘protective barrier’ to the phage. Differences in genetic diversity may have occurred at the start of treatment, with the patient infected by a mix of closely-related strains of the same bacterium, or between infection commencement and start of phage treatment. Unfortunately, we do not have estimates of genetic diversity over the course of infection as we attained no blood samples for comparison. However, it would be an interesting area of future research to examine how diversity in an infecting population affects phage therapy efficacy.

The probability of phage resistance evolving *in vivo* was greatly overestimated in both single and especially mixed isolate *in vitro* populations (41.6% replicates of mixed-isolate, 16.7% of single isolate replicates). In cultures where resistance emerged, this typically reached fixation with recovery of populations comparable to controls (implying minimal costs of resistance) and a lack of phage replication. This may be more likely to occur *in vitro* than *in vivo* as populations are only under stress from phage, without co-acting stressors found *in vivo* (Hernandez and Koskella 2019). Increased stress on bacteria populations can reduce population sizes and slows growth, reducing mutation supply rate for resistance (Gómez and Buckling 2011; Friman and Buckling 2013; Lopez Pascua et al. 2014; Harrison et al. 2015; Mumford and Friman 2017). Additionally, phage resistance may be constrained if it increases susceptibility to other stressors such as antibiotics (Chan et al. 2016), the immune system (Flyg et al. 1980; Roach et al. 2017), or reduces competitive ability with other bacteria (Winter et al. 2010). Importance of these trade-offs is illustrated in case studies where phage resistance emerges but does not result in treatment failure (Castledine et al. 2022; Van Nieuwenhuyse et al. 2022; Pirnay et al. 2024). The considerably greater rate of resistance evolution in mixed versus single populations can be simply explained by greater genetic diversity increases the evolutionary potential of genetically diverse populations (Brockhurst et al. 2003; Morgan et al. 2010).

These results show the challenges of making phage therapy predictions using *in vitro* experiments. Using mixed-isolates gives bacteria a relative advantage, giving possible insight into a ‘worse case scenario’ where phage have reduced efficacy and resistance may emerge. This may perhaps be useful in preparation for then selecting additional phage for cocktail development, with synergistic antibiotics which is the standard for phage therapy delivery (Pirnay et al. 2024). Additionally, these experiments may be beneficial in identifying conditions in which phage achieve best results, with interventions such as pus removal prior to phage therapy perhaps recommended to enhance treatment success. Results would need to be interpreted with understanding that bacteria *in vivo* would also be disadvantaged with the presence of the immune system and, depending on infection type, host microbiome. *In vitro* experiments may give more accurate predictions in relatively more simple infection scenarios, such as our prior work where *in vitro* experiments predicted resistance-virulence trade-offs in a relatively simple nasal colonisation scenario (Castledine et al. 2022). Although similar mutations were observed, different mechanisms of phage resistance were observed *in vivo* and *in vitro* which reinforces that *in vitro* environments shape evolutionary scenarios differentially.

Further work is needed to understand what variables shape bacteria-phage (co)evolution *in vivo* over the course of phage therapy. Work should also aim to integrate and analyse real phage therapy samples wherever possible, to give more accurate insights into real clinical scenarios. As this study focusses on a single phage therapy case study, generalisability is limited. However, all phage therapy cases undergo similar procedures in which phage efficacy is tested *in vitro* prior to clinical application. An interesting addition to published case reports would be to reflect on the predictability these *in vitro* studies had on clinical outcomes, with a clear statement on whether authors used single or multiple isolates to test phage efficacy. Testing phage resistance *in vitro* and *in vivo* in these assays would also provide a more comprehensive understanding behind treatment success and failure (Abedon, 2017).

## Acknowledgements

We thank Mzia Kutateladze, Director of the Eliava Institute, for providing the phage strain ISP.

