## Supplementary material for "Predicting clinical phage therapy outcomes *in vitro*: results using mixed versus single isolates from an MRSA case study": Figure S

**Supplementary Information**

Table S1. Details of samples from which bacteria and phage were successfully extracted. Y indicates successful extraction, with a pool of 24 isolates taken for analysis. N indicates no bacteria or phage detected in the sample.

| Sample | Day of treatment | Bacteria | Phage |
| --- | --- | --- | --- |
| Pleural fluid | 0 | Y | N |
| Pus | 0 | Y | N |
| Pleural fluid | 6 (pre-phage dose) | N | Y |
| Pleural fluid | 6 (post-phage dose) | N | Y |
| Bronchoalveolar lavage | 6 | Y | N |

Table S2. Tukey HSD comparisons comparing bacteria densities across cultures initiated from four different single isolates or a ‘mix’ of 24 isolates, in the presence (Y) and absence (N) of phage. P-values are adjusted by the tukey method for comparing a family of five estimates.

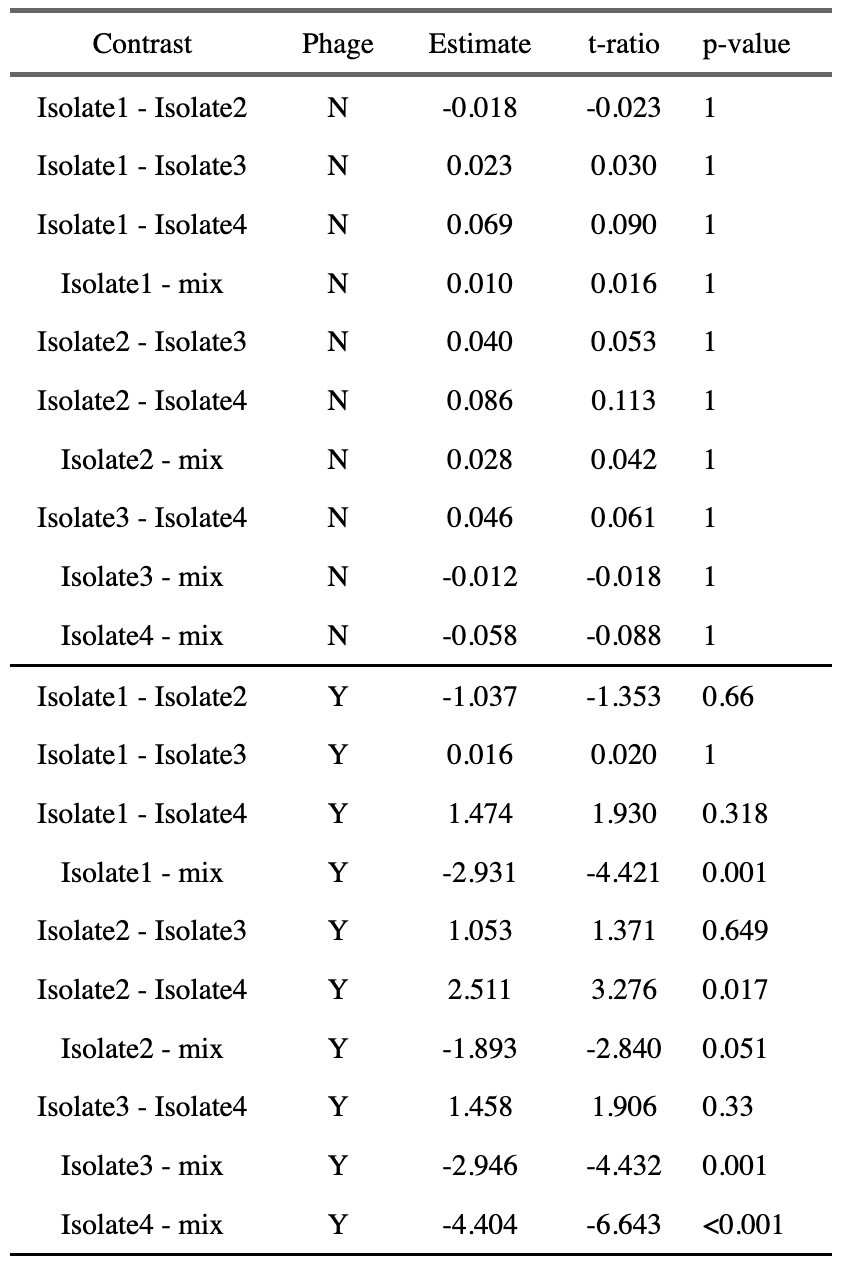

Table S3. Tukey HSD comparisons comparing bacteria densities through time in the presence (Y) and absence (N) of phage. Numbers under contrast indicate day of experiment bacteria densities were measured. P-values are adjusted by the tukey method for comparing a family of six estimates.

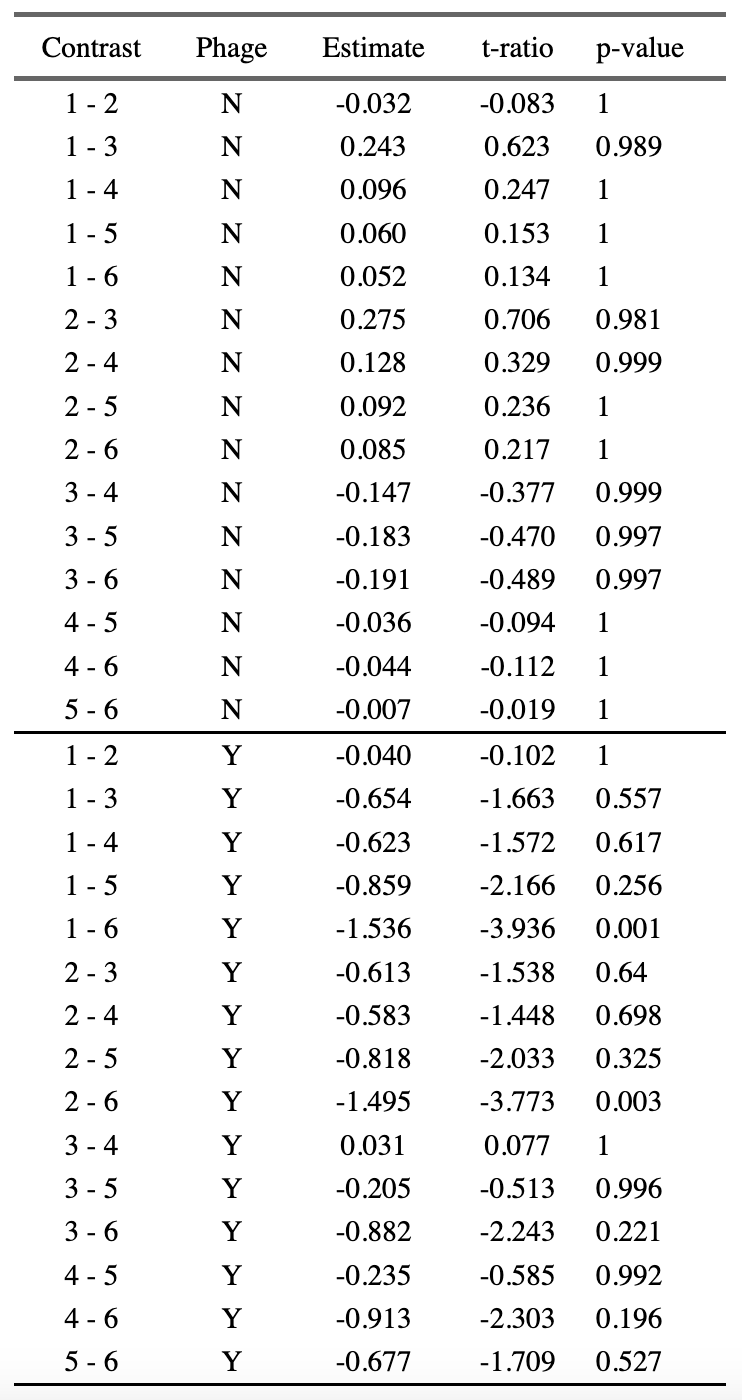

Table S4. One-sample t-tests comparing phage densities within each treatment, at each time-point, to the base-line inoculum density. Densities greater than this value indicate that phages are replicating. P-values adjusted for multiple testing using the false-discovery-rate (‘fdr’) method.

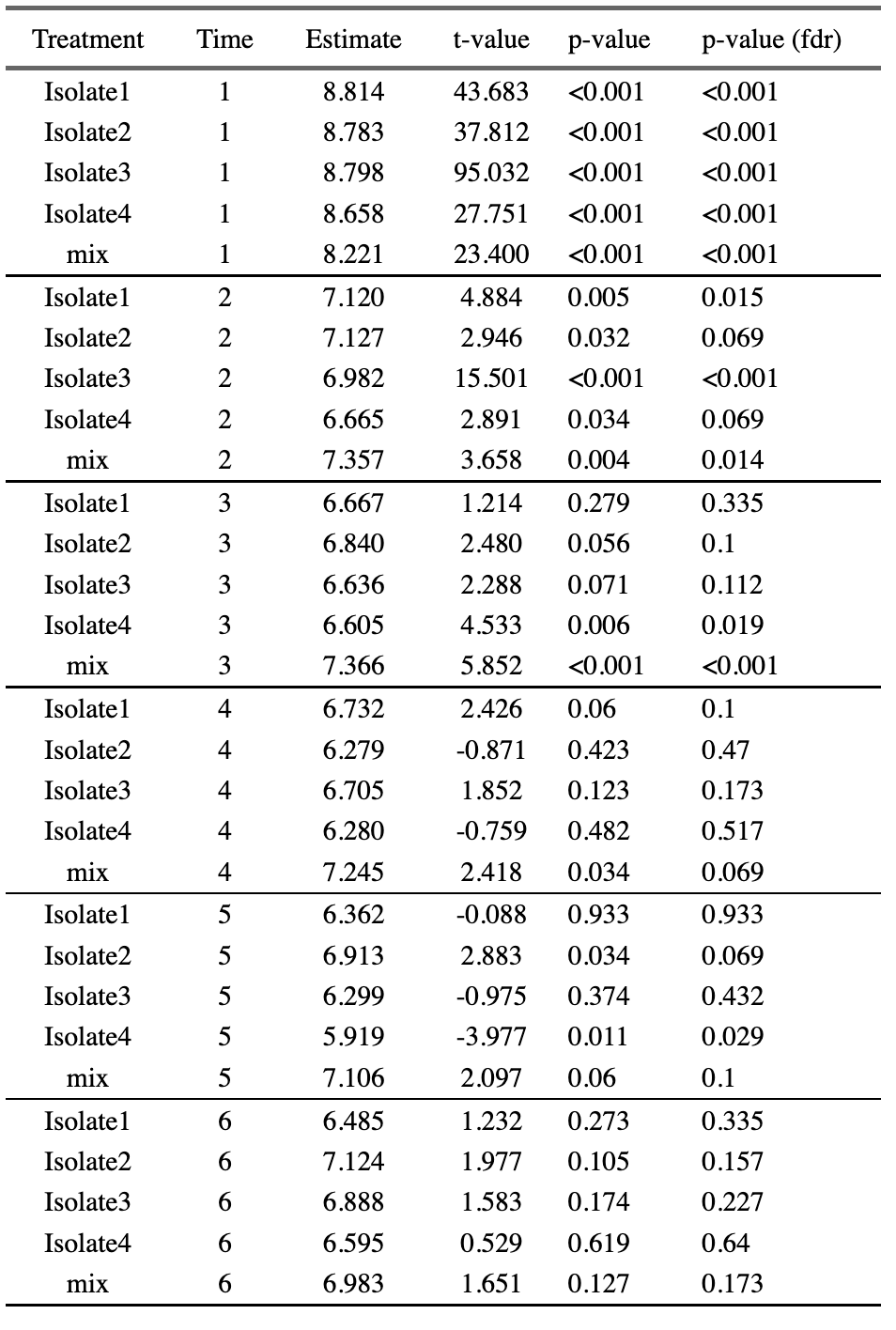
